# A Gaussian process model of human electrocorticographic data

**DOI:** 10.1101/121020

**Authors:** Lucy L. W. Owen, Tudor A. Muntianu, Andrew C. Heusser, Patrick Daly, Katherine Scangos, Jeremy R. Manning

## Abstract

We present a model-based method for inferring full-brain neural activity at millimeter-scale spatial resolutions and millisecond-scale temporal resolutions using standard human intracranial recordings. Our approach makes the simplifying assumptions that different people’s brains exhibit similar correlational structure, and that activity and correlation patterns vary smoothly over space. One can then ask, for an arbitrary individual’s brain: given recordings from a limited set of locations in that individual’s brain, along with the observed spatial correlations learned from other people’s recordings, how much can be inferred about ongoing activity at *other* locations throughout that individual’s brain? We show that our approach generalizes across people and tasks, thereby providing a person- and task-general means of inferring high spatiotemporal resolution full-brain neural dynamics from standard low-density intracranial recordings.

## Introduction

Modern human brain recording techniques are fraught with compromise (Sejnowski et al. 2014). Commonly used approaches include functional magnetic resonance imaging (fMRI), scalp electroencephalography (EEG), and magnetoencephalography (MEG). For each of these techniques, neuroscientists and electrophysiologists must choose to optimize spatial resolution at the cost of temporal resolution (e.g., as in fMRI) or temporal resolution at the cost of spatial resolution (e.g., as in EEG and MEG). A less widely used approach (due to requiring work with neurosurgical patients) is to record from electrodes implanted directly onto the cortical surface (electrocorticography; ECoG) or into deep brain structures (intracranial EEG; iEEG). However, these intracranial approaches also require compromise: the high spatiotemporal resolution of intracranial recordings comes at the cost of substantially reduced brain coverage, since safety considerations limit the number of electrodes one may implant in a given patient’s brain. Further, the locations of implanted electrodes are determined by clinical, rather than research, needs.

An increasingly popular approach is to improve the effective spatial resolution of MEG or scalp EEG data by using a geometric approach called *beamforming* to solve the biomagnetic or bioelectrical inverse problem (Sarvas 1987). This approach entails using detailed brain conductance models (often informed by high spatial resolution anatomical MRI images) along with the known sensor placements (localized precisely in 3D space) to reconstruct brain signals originating from theoretical point sources deep in the brain (and far from the sensors). Traditional beamforming approaches must overcome two obstacles. First, the inverse problem beamforming seeks to solve has infinitely many solutions. Researchers have made progress towards constraining the solution space by assuming that signal-generating sources are localized on a regularly spaced grid spanning the brain and that individual sources are small relative to their distances to the sensors (Baillet et al. 2001; Hillebrand et al. 2005; Snyder 1991). The second, and in some ways much more serious, obstacle is that the magnetic fields produced by external (noise) sources are substantially stronger than those produced by the neuronal changes being sought (i.e., at deep structures, as measured by sensors at the scalp). This means that obtaining adequate signal quality often requires averaging the measured responses over tens to hundreds of responses or trials (e.g., see review by Hillebrand et al. 2005).

Another approach to obtaining high spatiotemporal resolution neural data has been to collect fMRI and EEG data simultaneously. Simultaneous fMRI-EEG has the potential to balance the high spatial resolution of fMRI with the high temporal resolution of scalp EEG, thereby, in theory, providing the best of both worlds. In practice, however, the signal quality of both recordings suffers substantially when the two techniques are applied simultaneously (e.g., see review by Huster et al. 2012). In addition, the experimental designs that are ideally suited to each technique individually are somewhat at odds. For example, fMRI experiments often lock stimulus presentation events to the regularly spaced image acquisition time (TR), which maximizes the number of post-stimulus samples. By contrast, EEG experiments typically employ jittered stimulus presentation times to maximize the experimentalist’s ability to distinguish electrical brain activity from external noise sources such as from 60 Hz alternating current power sources.

The current “gold standard” for precisely localizing signals and sampling at high temporal resolution is to take (ECoG or iEEG) recordings from implanted electrodes (but from a limited set of locations in any given brain). This begs the following question: what can we infer about the activity exhibited by the rest of a person’s brain, given what we learn from the limited intracranial recordings we have from their brain and additional recordings taken from *other* people’s brains? Here we develop an approach, which we call *SuperEEG*^1^, based on Gaussian process regression (Rasmussen 2006). SuperEEG entails using data from multiple people to estimate activity patterns at arbitrary locations in each person’s brain (i.e., independent of their electrode placements). We test our SuperEEG approach using two large datasets of intracranial recordings (Ezzyat et al. 2017, 2018; Horak et al. 2017; Kragel et al. 2017; Kucewicz et al. 2017, 2018; Lin et al. 2017; Manning et al. 2011, 2012; Sederberg et al. 2003, 2007a,b; Solomon et al. 2018; Weidemann et al. 2019). We show that the SuperEEG algorithm recovers signals well from electrodes that were held out of the training dataset. We also examine the factors that influence how accurately activity may be estimated (recovered), which may have implications for electrode design and placement in neurosurgical applications.

## Approach

The SuperEEG approach to inferring high temporal resolution full-brain activity patterns is outlined and summarized in Figure 1. We describe (in this section) and evaluate (in *Results*) our approach using two large previously collected datasets comprising multi-session intracranial recordings. Dataset 1 comprises multi-session recordings taken from 6876 electrodes implanted in the brains of 88 epilepsy patients (Manning et al. 2011, 2012; Sederberg et al. 2003, 2007a,b). Each recording session lasted from 0.2–3 h (total recording time: 0.3–14.2 h; Fig. S6E). During each recording session, the patients participated in a free recall list learning task, which lasted for up to approximately 1 h. In addition, the recordings included “buffer” time (the length varied by patient) before and after each experimental session, during which the patients went about their regular hospital activities (confined to their hospital room, and primarily in bed). These additional activities included interactions with medical staff and family, watching television, reading, and other similar activities. For the purposes of the Dataset 1 analyses presented here, we aggregated all data across each recording session, including recordings taken during the main experimental task as well as during non-experimental time. We used Dataset 1 to develop our main SuperEEG approach, and to examine the extent to which SuperEEG might be able to generate task-general predictions. Dataset 2 comprised multi-session recordings from 14860 electrodes implanted in the brains of 131 epilepsy patients (Ezzyat et al. 2017, 2018; Horak et al. 2017; Kragel et al. 2017; Kucewicz et al. 2017, 2018; Lin et al. 2017; Solomon et al. 2018; Weidemann et al. 2019). Each recording session lasted from 0.4–2.2 h (total recording time: 0.4–6.6 h; Fig. S6K). Whereas Dataset 1 included recordings taken as the patients participated in a variety of activities, Dataset 2 included recordings taken as each patient performed each of two specific experimental memory tasks: a random word list free recall task (Experiment 1) and a categorized word list free recall task (Experiment 2). We used Dataset 2 to further examine the ability of SuperEEG to generalize its predictions within versus across tasks. Figure S6 provides additional information about both datasets.

**Figure 1:**
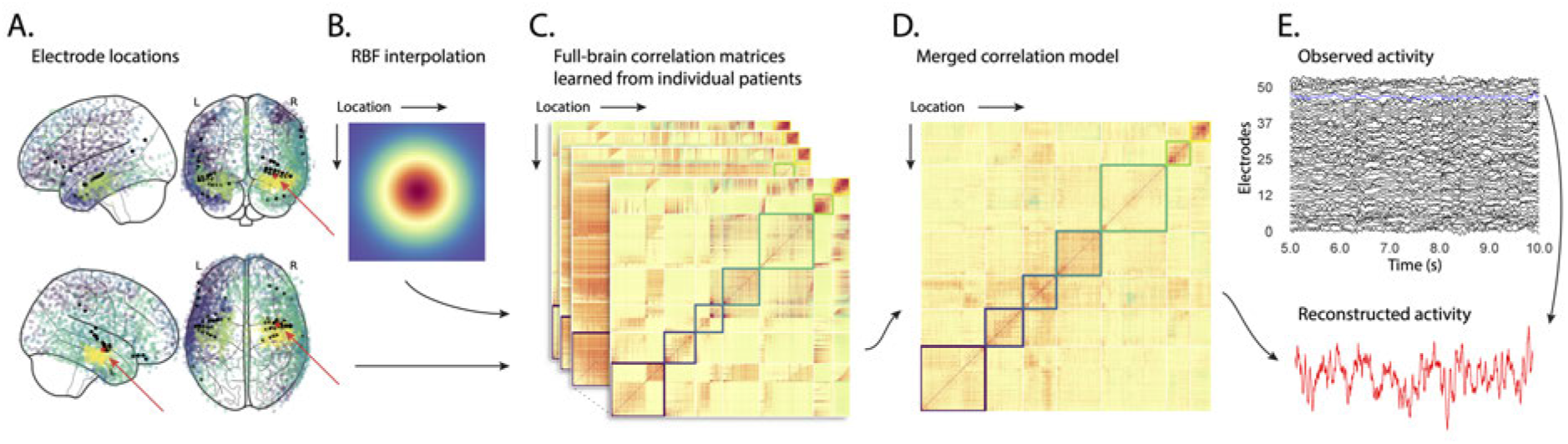
Methods overview. **A. Electrode locations.** Each dot reflects the location of a single electrode implanted in the brain of a Dataset 1 patient. A held-out recording location from one patient is indicated in red, and the patient’s remaining electrodes are indicated in black. The electrodes from the remaining patients are colored by *k*-means cluster (computed using the full-brain correlation model shown in Panel D). **B. Radial basis function kernel.** Each electrode contributed by the patient (black) weights on the full set of locations under consideration (all dots in Panel A, defined as 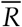 in the text). The weights fall off with positional distance (in MNI152 space) according to an RBF. **C. Per-patient correlation matrices.** After computing the pairwise correlations between the recordings from each patient’s electrodes, we use RBF-weighted averages to estimate correlations between all locations in 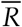. We obtain an estimated full-brain correlation matrix using each patient’s data. **D. Merged correlation model.** We combine the per-patient correlation matrices (Panel C) to obtain a single full-brain correlation model that captures information contributed by every patient. Here we have sorted the rows and columns to reflect *k*-means clustering labels (using *k*=7; Yeo et al. 2011), whereby we grouped locations based on their correlations with the rest of the brain (i.e., rows of the matrix displayed in the panel). The boundaries denote the cluster groups. The rows and columns of Panel C have been sorted using the Panel D-derived cluster labels. **E. Reconstructing activity throughout the brain.** Given the observed recordings from the given patient (shown in black; held-out recording is shown in blue), along with a full-brain correlation model (Panel D), we use Equation 12 to reconstruct the most probable activity at the held-out location (red).

We first applied fourth order Butterworth notch filters to remove 60 Hz (± 0.5 Hz) line noise from every recording (from every electrode). Next, we downsampled the recordings (regardless of the original samplerate) to 250 Hz. This downsampling step served to both normalize for differences in sampling rates across patients and to ease the computational burden of our subsequent analyses. We then excluded any electrodes that showed putative epileptiform activity. Specifically, we excluded from further analysis any electrode that exhibited a maximum kurtosis of 10 or greater across all of that patient’s recording sessions. We also excluded any patients with fewer than 2 electrodes that passed this criteria, as the SuperEEG algorithm requires measuring correlations between 2 or more electrodes from each patient. For Dataset 1, this yielded clean recordings from 4168 electrodes implanted throughout the brains of 67 patients (Fig. 1A, colored dots); for Dataset 2, this yielded clean recordings from 5023 electrodes implanted in 78 patients. Each individual patient contributed electrodes from a limited set of brain locations, which we localized in a common space (MNI152; Grabner et al. 2006); an example Dataset 1 patient’s 54 electrodes that survived the kurtosis thresholding procedure are highlighted in black and red (Fig. 1A).

The recording from a given electrode is maximally informative about the activity of the neural tissue immediately surrounding its recording surface. However, brain regions that are distant from the recording surface of the electrode also contribute to the recording, albeit (ceteris paribus) to a much lesser extent. One mechanism underlying these contributions is volume conduction. The precise rate of falloff due to volume conduction (i.e., how much a small volume of brain tissue at location *x* contributes to the recording from an electrode at location η) depends on the size of the recording surface, the electrode’s impedance, and the conductance profile of the volume of brain between *x* and *η*. As an approximation of this intuition, we place a Gaussian radial basis function (RBF) at the location *η* of each electrode’s recording surface (Fig. 1B). We use the values of the RBF at any brain location *x* as a rough estimate of how much structures around *x* contributed to the recording from location *η*:

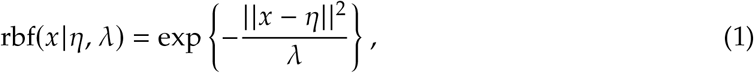

where the width variable λ is a parameter of the algorithm (which may in principle be set according to location-specific tissue conductance profiles) that governs the level of spatial smoothing. In choosing λ for the analyses presented here, we sought to maximize spatial resolution (which implies a small value of λ) while also maximizing the algorithm’s ability to generalize to any location throughout the brain, including those without dense electrode coverage (which implies a large value of λ). Here we set λ = 20, guided in part by our prior related work (Manning et al. 2014, 2018), and in part by examining the brain coverage with non-zero weights achieved by placing RBFs at each electrode location in Dataset 1 and taking the sum (across all electrodes) at each voxel in a 4 mm^3^ MNI brain. (We then held λ fixed for our analyses of Dataset 2.) We note that this value could in theory be further optimized, e.g., using cross validation or a formal model (e.g., Manning et al. 2018).

A second mechanism whereby a given region *x* can contribute to the recording at *η* is through (direct or indirect) anatomical connections between structures near *x* and *η*. Although anatomical and functional correlations can differ markedly (e.g., Adachi et al. 2012; Goñi et al. 2014; Honey et al. 2009), we use temporal correlations in the data to estimate these anatomical connections (Becker et al. 2018). Let 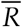 be the set of locations at which we wish to estimate local field potentials, and let 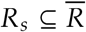 be set of locations at which we observe local field potentials from patient *s* (excluding the electrodes that did not pass the kurtosis test described above). In the analyses below we define 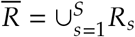. We can calculate the expected inter-electrode correlation matrix for patient *s*, where *C_s,k_* (*i*, *j*) is the correlation between the time series of voltages for electrodes *i* and *j* from subject *s* during session *k*, using:

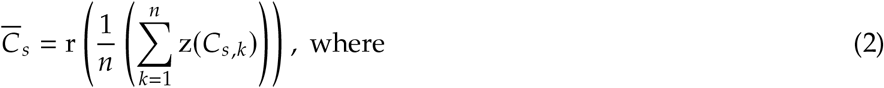

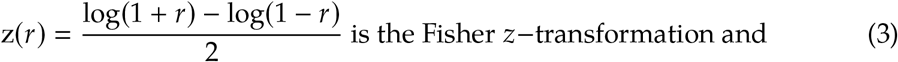

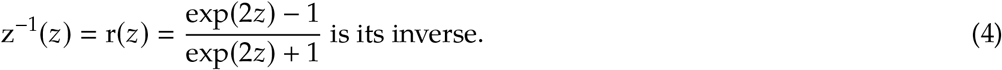

Next, we use Equation 1 to construct a number of to-be-estimated locations by number of patient electrode locations weight matrix, *W*_*s*_. Specifically, *W*_*s*_ approximates how informative the recordings at each location in *R*_*s*_ are in reconstructing activity at each location in 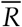, where the contributions fall off with an RBF according to the distances between the corresponding locations:

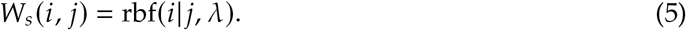

Given this weight matrix, *W*_*s*_, and the observed inter-electrode correlation matrix for patient *s*, 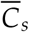, we can estimate the correlation matrix for all locations in 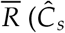; Fig. 1C) using:

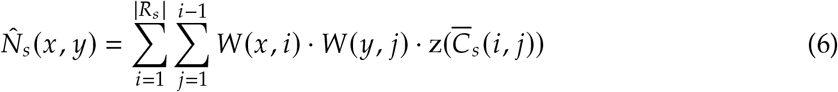

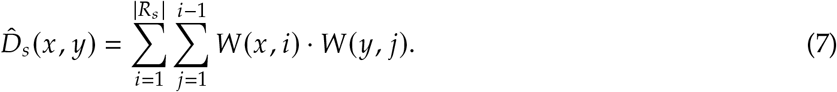

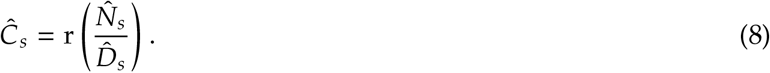

After estimating the numerator 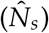 and denominator 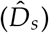 placeholders for each 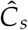, we aggregate these estimates across the *S* patients to obtain a single expected full-brain correlation matrix (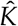; Fig. 1D):

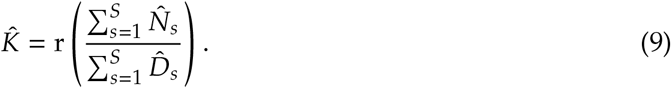

Intuitively, the numerators capture the general structures of the patient-specific estimates of full-brain correlations, and the denominators account for which locations were near the implanted electrodes in each patient. To obtain 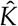, we compute a weighted average across the estimated patient-specific full-brain correlation matrices, where patients with observed electrodes near a particular set of locations in 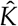 contribute more to the estimate.

Having used the multi-patient data to estimate a full-brain correlation matrix at the set of locations in 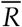 that we wish to know about, we next use 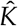 to estimate activity patterns everywhere in 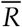, given observations at only a subset of locations in 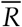 (Fig. 1E).

Let *α*_*s*_ be the set of indices of patient *s*’s electrode locations in 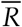 (i.e., the locations in *R*_*s*_), and let *β*_*s*_ be the set of indices of all other locations in 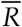. In other words, *β*_*s*_ reflects the locations in 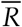 where we did not observe a recording for patient *s* (these are the recording locations we will want to fill in using SuperEEG). We can sub-divide 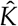 as follows:

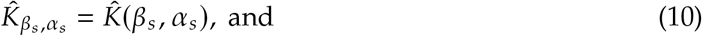

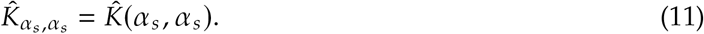

Here 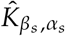 represents the correlations between the “unknown” activity at the locations indexed by *β*_*s*_ and the observed activity at the locations indexed by *α*_*s*_, and 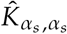 represents the correlations between the observed recordings (at the locations indexed by *α*_*s*_).

Let 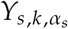 be the number-of-timepoints (*T*) by |*α*_*s*_| matrix of (observed) voltages from the electrodes in *α*_*s*_ during session *k* from patient *s*. Then we can estimate the voltage from patient *s*’*s k^th^* session at the locations in *β*_*s*_ as follows (Rasmussen 2006):

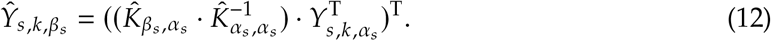

This equation is the foundation of the SuperEEG algorithm. Whereas we observe recordings only at the locations indexed by *α*_*s*_, Equation 12 allows us to estimate the recordings at all locations indexed by *β*_*s*_, which we can define *a priori* to include any locations we wish, throughout the brain. This yields estimates of the time-varying voltages at *every* location in 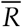, provided that we define 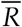 in advance to include the union of all of the locations in *R*_*s*_ and all of the locations at which we wish to estimate recordings (i.e., a timeseries of voltages).

We designed our approach to be agnostic to electrode impedances, as electrodes that do not exist do not have impedances. Therefore our algorithm recovers voltages in standard deviation (*z*-scored) units rather than attempting to recover absolute voltages. (This property reflects the fact that 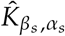 and 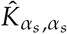 are correlation matrices rather than covariance matrices.) Also, we note that Equation 12 requires computing a *T* by *T* matrix, which can become computationally expensive when *T* is very large (e.g., for the Dataset 1 patient with the longest recording time, *T* = 12, 786, 750; also see Fig. S6, Panels E and K). However, because Equation 12 is time invariant, we may compute 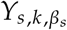 in a piecewise manner by filling in 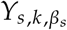 one row at a time (using the corresponding samples from 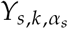).

The SuperEEG algorithm described above and in Figure 1 allows us to estimate, up to a constant scaling factor, local field potentials (LFPs) for each patient at all arbitrarily chosen locations in the set 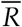, *even if we did not record that patient’s brain at all of those locations*. We next turn to an evaluation of the accuracy of those estimates.

## Results

We used a cross-validation approach to test the accuracy with which the SuperEEG algorithm reconstructs activity throughout the brain. For each patient in turn, we estimated full-brain correlation matrices (Eqn. 9) using data from all of the *other* patients. This step ensured that the data we were reconstructing could not also be used to estimate the between-location correlations that drove the reconstructions via Equation 12 (otherwise the analysis would be circular). For that held-out patient, we held out each electrode in turn. We used Equation 12 to reconstruct activity at the held-out electrode location, using the correlation matrix learned from all other patients’ data as 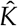, and using activity recorded from the other electrodes from the held-out patient as 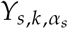. (For analyses examining the stability of our estimates of 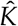 across time and patients, see Figs. S7 and S8, respectively). We then asked: how closely did each of the SuperEEG-estimated recordings at those electrodes match the observed recordings from those electrodes (i.e., how closely did the estimated 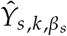 match the observed 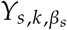)?

We used this general approach to quantify the algorithm’s performance across the full dataset. For each held-out electrode, from each held-out patient in turn, we computed the average correlation (across recording sessions) between the SuperEEG-reconstructed voltage traces and the observed voltage traces from that electrode. For this analysis we set 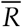 to be the union of all electrode locations across all patients. This yielded a single correlation coefficient for each electrode location in 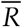, reflecting how well the SuperEEG algorithm was able to recover the recording at that location by incorporating data across patients (black histogram in Fig. 2A, map in Fig. 2C). The observed distribution of correlations was centered well above zero (mean: *r* = 0.51; *t*-test comparing mean of distribution of *z*-transformed average patient correlation coefficients to 0: *t*(66) = 23.55, *p* < 10^−10^), indicating that the SuperEEG algorithm recovers held-out activity patterns substantially better than random guessing.

**Figure 2:**
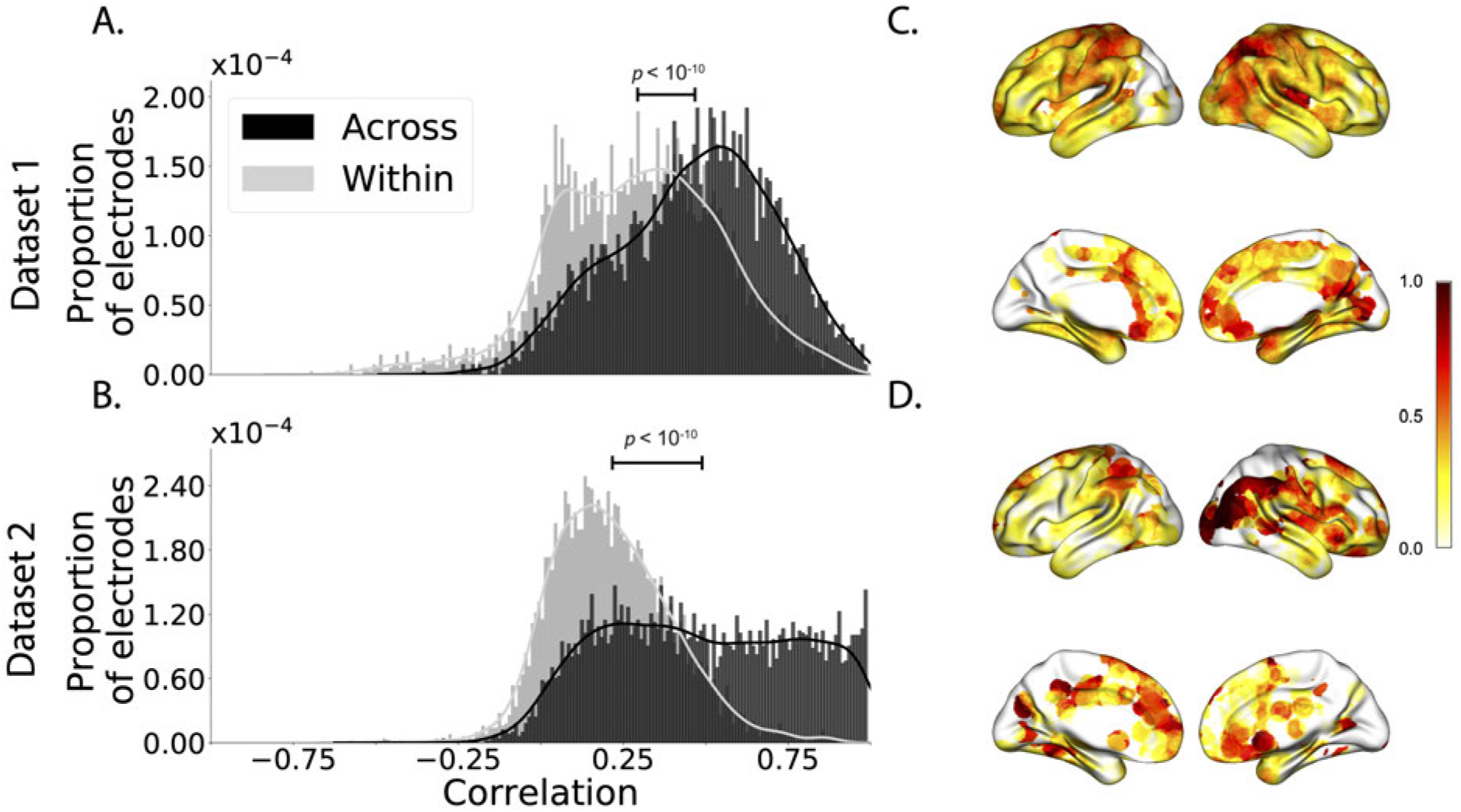
Reconstruction accuracy across all electrodes in two ECoG datasets. **A. Distributions of correlations between observed versus reconstructed activity by electrode, for Dataset 1.** The across-patient distribution (black) reflects reconstruction accuracy (correlation) using a correlation model learned from all but one patient’s data, and then applied to that held-out patient’s data. The within-patient distribution (gray) reflects performance using a correlation model learned from the same patient who contributed the to-be-reconstructed electrode. **B. Distributions of correlations for Dataset 2.** This panel is in the same format as Panel A, but reflects results obtained from Dataset 2. The histograms aggregate data across both Dataset 2 experiments; for results broken down by experiment see Figures S2 and S3. **C.–D. Reconstruction accuracy by location.** The colors denote the average across-session correlations, using the across-patient correlation model, between the observed and reconstructed activity at the given electrode location projected to the cortical surface (Combrisson et al. 2019). Panel C displays the map for Dataset 1 and Panel D displays the map for Dataset 2.

Next, we compared the quality of these across-participant reconstructions (i.e., computed using a correlation model learned from other patients’ data) to reconstructions generated using a correlation model trained using the in-patient’s data. In other words, for this within-patient benchmark analysis we estimated 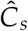 (Eqn. 8) for each patient in turn, using recordings from all of that patient’s electrodes except at the location we were reconstructing. These within-patient reconstructions serve as an estimate of how well data from all of the other electrodes from that single patient may be used to estimate held-out data from the same patient. This allows us to ask how much information about the activity at a given electrode might be inferred through (a) volume conductance or other sources of “leakage” from activity patterns measured from the patient’s other electrodes and (b) across-electrode correlations learned from that single patient. As shown in Figure 2A (gray histogram), the distribution of within-patient correlations was centered well above zero (mean: *r* = 0.32; *t*-test comparing mean of distribution of *z*-transformed average patient correlation coefficients to 0: *t*(66) = 15.16, *p* < 10^−10^). However, the across-patient correlations were substantially higher (*t*-test comparing average *z*-transformed within versus across patient electrode correlations: *t*(66) = 9.17, *p* < 10^−10^). This is an especially conservative test, given that the across-patient SuperEEG reconstructions exclude (from the correlation matrix estimates) all data from the patient whose data is being reconstructed. We repeated each of these analyses on a second independent dataset and found similar results (Fig. 2B, D; within versus across reconstruction accuracy: *t*(77) = 11.25, *p* < 10^−10^). We also replicated this result separately for each of the two experiments from Dataset 2 (Fig. S3). This overall finding, that reconstructions of held-out data using correlation models learned from *other* patient’s data yield higher reconstruction accuracy than correlation models learned from the patient whose data is being reconstructed, has two important implications. First, it implies that distant electrodes provide additional predictive power to the data reconstructions beyond the information contained solely in nearby electrodes. This follows from the fact that each patient’s grid, strip, and depth electrodes are implanted in a unique set of locations, so for any given electrode the closest electrodes in the full dataset tend to come from the same patient. Second, it implies that the spatial correlations learned using the SuperEEG algorithm are, to some extent, similar across people.

The recordings we analyzed from Dataset 1 comprised data collected as the patients performed a variety of (largely idiosyncratic) tasks throughout each day’s recording session. That we observed reliable reconstructions across patients suggests that the spatial correlations derived from the SuperEEG algorithm are, to some extent, similar across tasks. We tested this finding more directly using Dataset 2. In Dataset 2, the recordings were limited to times when each patient was participating in one of two experiments. Experiment 1 is a random-word list free recall task; Experiment 2 is a categorized list free recall task (24 patients participated in both). We wondered whether a correlation model learned from data from one experiment might yield good predictions of data from the other experiment. Further, we wondered about the extent to which it might be beneficial or harmful to combine data across tasks.

To test the task-specificity of the SuperEEG-derived correlation models, we restricted the dataset to the 24 patients that participated in both experiments and repeated the above within- and across-patient cross validation procedures separately for Experiment 1 and Experiment 2 data from Dataset 2. We then compared the reconstruction accuracies for held-out electrodes, for models trained within versus across the two experiments, or combining across both experiments (Fig. S1). In every case we found that across-patient models trained using data from all other patients out-performed within-patient models trained on data only from the subject contributing the given electrode (*t*s(23) > 6.50, *p*s< 10^−5^). All reconstruction accuracies also reliably exceeded chance performance (*t*s(23) > 8.00, *p*s< 10^−8^). Average reconstruction accuracy was highest for the across-patient models limited to data from the same experiment (mean accuracy: *r* = 0.68); next-highest for the models that combined data across both experiments (mean accuracy: *r* = 0.61); and lowest for models trained across tasks (mean accuracy: *r* = 0.47). This pattern of results also held for each of the Dataset 2 experiments individually (Fig. S2). Taken together, these results indicate that there are reliable commonalities in the spatial correlations of full-brain activity across tasks, but that there are also reliable differences in these spatial correlations across tasks. Whereas reconstruction accuracy benefits from incorporating data from other patients, reconstruction accuracy is highest when constrained to within-task data, or data that includes a variety of tasks (e.g., Dataset 1, or combining across the two Dataset 2 experiments).

Although both datasets we examined provide good full-brain coverage (when considering data from every patient), electrodes were not sampled uniformly throughout the brain. For example, in our patient population, electrodes are more likely to be implanted in regions like the medial temporal lobe (MTL), and are rarely implanted in occipital cortex (Fig. 3A, B). Separately for each dataset, for each voxel in the 4 mm^3^ voxel MNI152 brain, we computed the proportion of electrodes in the dataset that were contained within a 20 MNI unit radius sphere centered on that voxel. We defined the *density* at that location as this proportion. Across Datasets 1 and 2, the electrode placement densities were similar (correlation by voxel: *r* = 0.6, *p* < 10^−10^). We wondered whether regions with good coverage might be associated with better reconstruction accuracy. For example, Figures 2C and D indicate that some electrodes in the MTL (which tends to be relatively densely sampled) have relatively high reconstruction accuracy, and occipital electrodes (which tends to be relatively sparsely sampled) tend to have relatively low reconstruction accuracy. To test whether this held more generally across the entire brain, for each dataset we computed the electrode placement density for each electrode from each patient (using the proportion of *other* patients’ electrodes within 20 MNI units of the given electrode). We then correlated these density values with the across-patient reconstruction accuracies for each electrode. We found no reliable correlation between reconstruction accuracy and density for Dataset 1 (*r* = 0.05, *p* = 0.70) and a reliable negative correlation for Dataset 2 (*r* = −0.21, *p* = 0.05). This suggests that the reconstruction accuracies we observed are *not* driven solely by sampling density, but rather may also reflect higher order properties of neural dynamics such as functional correlations between distant voxels (Betzel et al. 2017).

**Figure 3:**
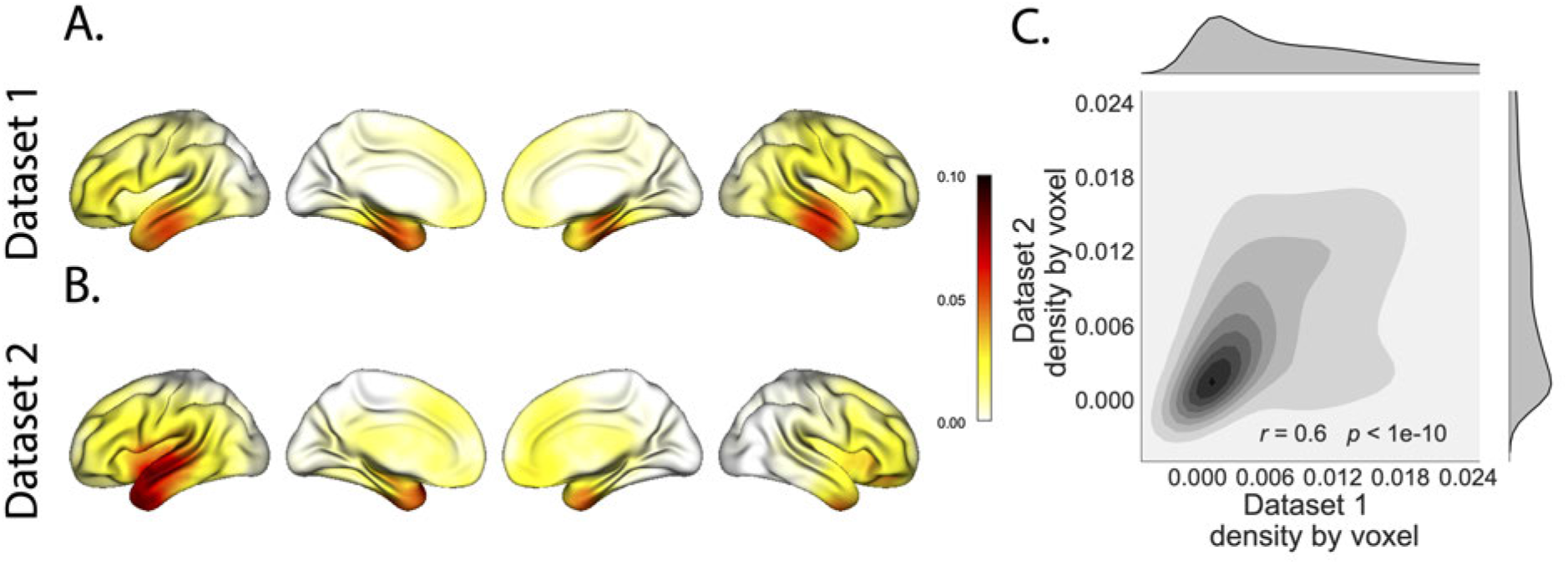
Electrode sampling density by location. **A. Electrode sampling density by voxel location in Dataset 1.** Each voxel is colored by the proportion of total electrodes in the dataset that are located within 20 MNI units of the given voxel. **B. Electrode sampling density by voxel location in Dataset 2.** This panel displays the sampling density map for Dataset 2, in the same format as Panel A. **C. Correspondence in sampling density by voxel location across Datasets 1 and 2.** The two-dimensional histogram displays the per-voxel sampling densities in the two Datasets, and the one-dimensional histograms display the proportions of voxels in each dataset with the given density value. The correlation reported in the panel is across voxels in the 4 mm^3^ MNI152 brain.

Prior work in humans and animals has shown that the spatial profile of the local field potential differs by frequency band (e.g., with respect to volume conductance properties and contribution to the local field potential; Buzsaki et al. 2012; Crone et al. 2011; Fries et al. 2007). For example, lower frequency components of the local field potential tend to have higher power and extend further in space than high-frequency components (e.g., Manning et al. 2009; Miller et al. 2007). We wondered whether the reconstructions we observed might be differently weighting or considering the contributions of activity at different frequency bands. We therefore examined a range of frequency bands (*δ*: 2–4 Hz; *θ*: 4–8 Hz; *α*: 8–12 Hz; *β*: 12–30 Hz; *γL* : 30–60 Hz; and *γH* : 60–100 Hz), along with a measure of broadband (BB) power. We used second-order Butterworth bandpass filters to compute the activity patterns within each narrow frequency band. We defined broadband power as the mean height of a linear robust regression fit in log-log space to the order 4 Morelet wavelet-computed power spectrum at 50 log-spaced frequencies from from 2–100 Hz (Manning et al. 2009). We then repeated our within-subject and across-subject cross-validated reconstruction accuracy tests (analogous to Fig. 2) separately for each frequency band (Fig. 4). (We also carried out a similar analysis on the Hilbert transform-computed spectral power within each narrow band; see Fig. S4.) Across both datasets, we found that our approach is best at reconstructing patterns of broadband activity (right-most bars in Figs. 4A and D), a correlate of population firing rate (Manning et al. 2009). We also achieved good reconstruction accuracy within each narrow frequency band (Figs. 4 and S4). Activity at lower frequencies (*δ*, *θ*, *α*, and *β*) tended to be reconstructed better than high-frequency patterns (*γL* and *γH*), with reconstruction accuracy peaking in the *θ* band. Overall, these results indicate that our approach is able to accurately recover information within the 2–100 Hz range.

**Figure 4:**
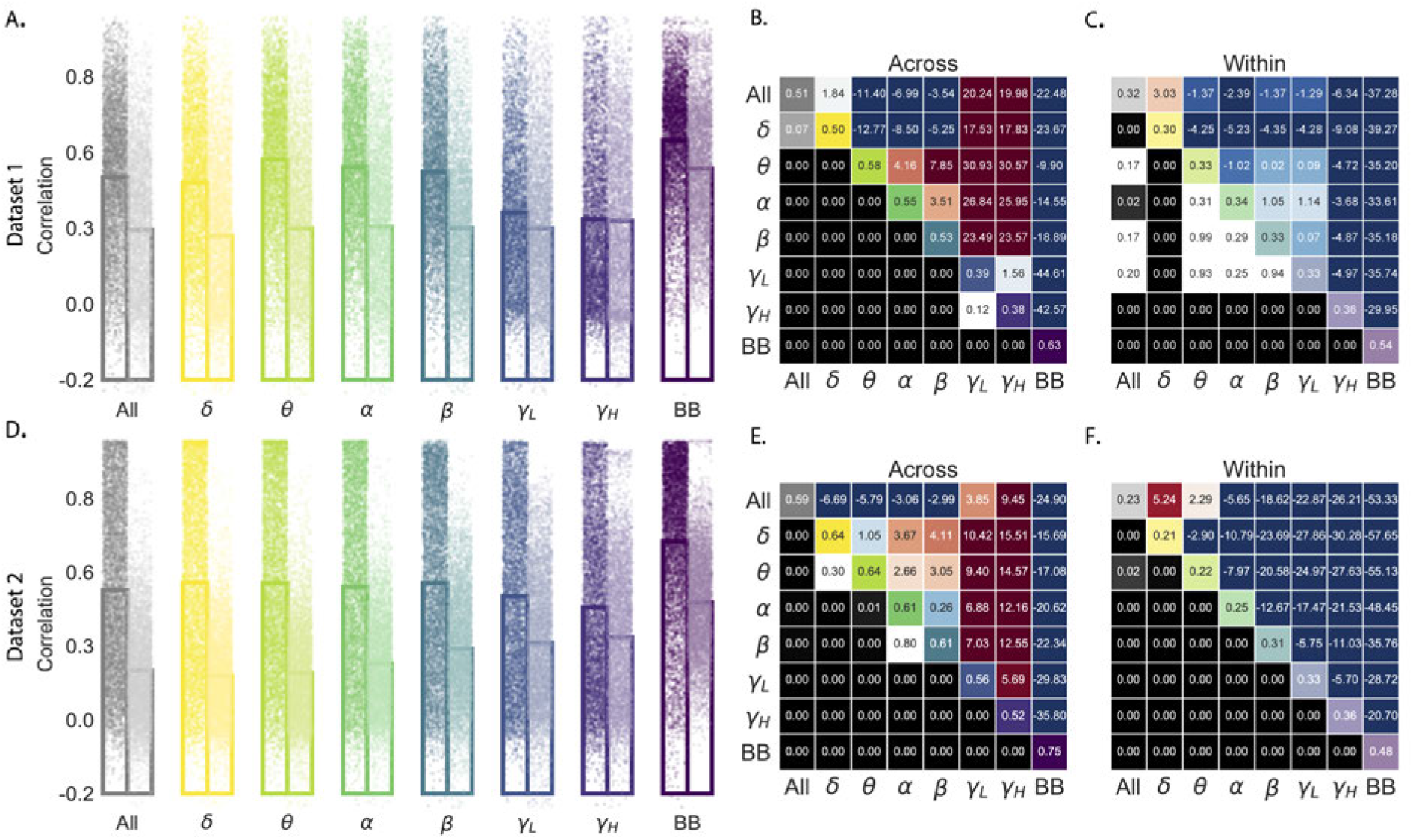
Reconstruction accuracy across all electrodes in two ECoG datasets for each frequency band. **A. Distributions of correlations between observed versus reconstructed activity by electrode for each frequency band in Dataset 1.** Each color denotes a different frequency band. Within each color group, the darker dots and bar on the left display the distribution (and mean) across-patient reconstruction accuracies (analogous to the black histograms in Fig. 2). The lighter dots and bar on the right display the distribution (and mean) within-patient reconstruction accuracies (analogous to the gray histograms in Fig. 2). Each dot indicates the reconstruction accuracy for one electrode in the dataset. To facilitate visual comparison with the frequency-specific results, the leftmost bars (gray) re-plot the histograms in Figure 2A. **B. Statistical summary of across-patient reconstruction accuracy by electrode for each frequency band in Dataset 1.** In the upper triangles of each map, warmer colors (positive *t*-values) indicate that the reconstruction accuracy for the frequency band in the given row was greater (via a two-tailed paired-sample *t*-test) than for the frequency band in the given column. Cooler colors (negative *t*-values) indicate that reconstruction accuracy for the frequency band in the given row was lower than for the frequency band in the given column. The lower triangles of each map denote the corresponding *p*-values for the *t*-tests. The diagonal entries display the average reconstruction accuracy within each frequency band. **C. Statistical summary of within-patient reconstruction accuracy by electrode for each frequency band in Dataset 1.** This panel displays the within-patient statistical summary, in the same format as Panel B. **D. Distributions of correlations between observed versus reconstructed activity by electrode, for each frequency band in Dataset 2.** This panel displays reconstruction accuracy distributions for each frequency band for Dataset 2. **E.–F. Statistical summaries of across-patient and within-patient reconstruction accuracy by electrode for each frequency band in Dataset 2.** These panels are in the same as Panels B and C, but display results from Dataset 2.

A basic assumption of our approach (and of most prior ECoG work) is that electrode recordings are most informative about the neural activity near the recording surface of the electrode. But if we consider that activity patterns throughout the brain are meaningfully correlated, are there particular implantation locations that, if recorded from a given patient’s brain, yield especially high reconstruction accuracies throughout the rest of their brain? For example, one might hypothesize that brain structures that are heavily interconnected with many other structures could be more informative about full-brain activity patterns than comparatively isolated structures. To test this hypothesis, we computed the average reconstruction accuracy across all of each patient’s electrodes (using our across-patients cross validation test; black histograms in Fig. 2A and B). We first labeled each patient’s electrodes, in each dataset, with the average reconstruction accuracy for that patient. In other words, we assigned every electrode from each patient the same value, reflecting how well the activity patterns for that patient were reconstructed. Next, for each voxel in the 4 mm^3^ MNI brain, we computed the average value across any electrode (from any patient) that came within 20 MNI units of that voxel’s center. This yielded an *information score* for each voxel, reflecting the (weighted) average reconstruction accuracy across any patients with electrodes near each voxel, where the averages were weighted to reflect patients who had more electrodes implanted near that location. We created a single map of these information scores for each dataset, highlighting regions that are especially informative about activity in *other* brain areas (Figs. 5A and B). Despite task and patient differences across the two datasets, we nonetheless found that the information score maps from both datasets were correlated (voxelwise correlation between information scores across the two datasets: *r* = 0.18, *p* < 10^−10^). Our finding that there were some commonalities between the two datasets’ information score maps lends support to the notion that different brain areas are (reliably) differently informative about full-brain activity patterns. We also examined the intersection between the top 10% most informative voxels across the two datasets (gray areas in Fig. 5C, networks shown in Fig. 6A, top row). Supporting the notion that structures that are highly interconnected with the rest of the brain are most informative about full-brain activity patterns, the intersecting set of voxels with the highest information scores included major portions of the dorsal attention network (e.g., inferior parietal lobule, precuneus, inferior temporal gyrus, thalamus, and striatum) as well as some portions of the default mode network (e.g., angular gyrus) that are highly interconnected with a large proportion of the brain’s gray matter (e.g., Tomasi and Volkow 2011).

**Figure 5:**
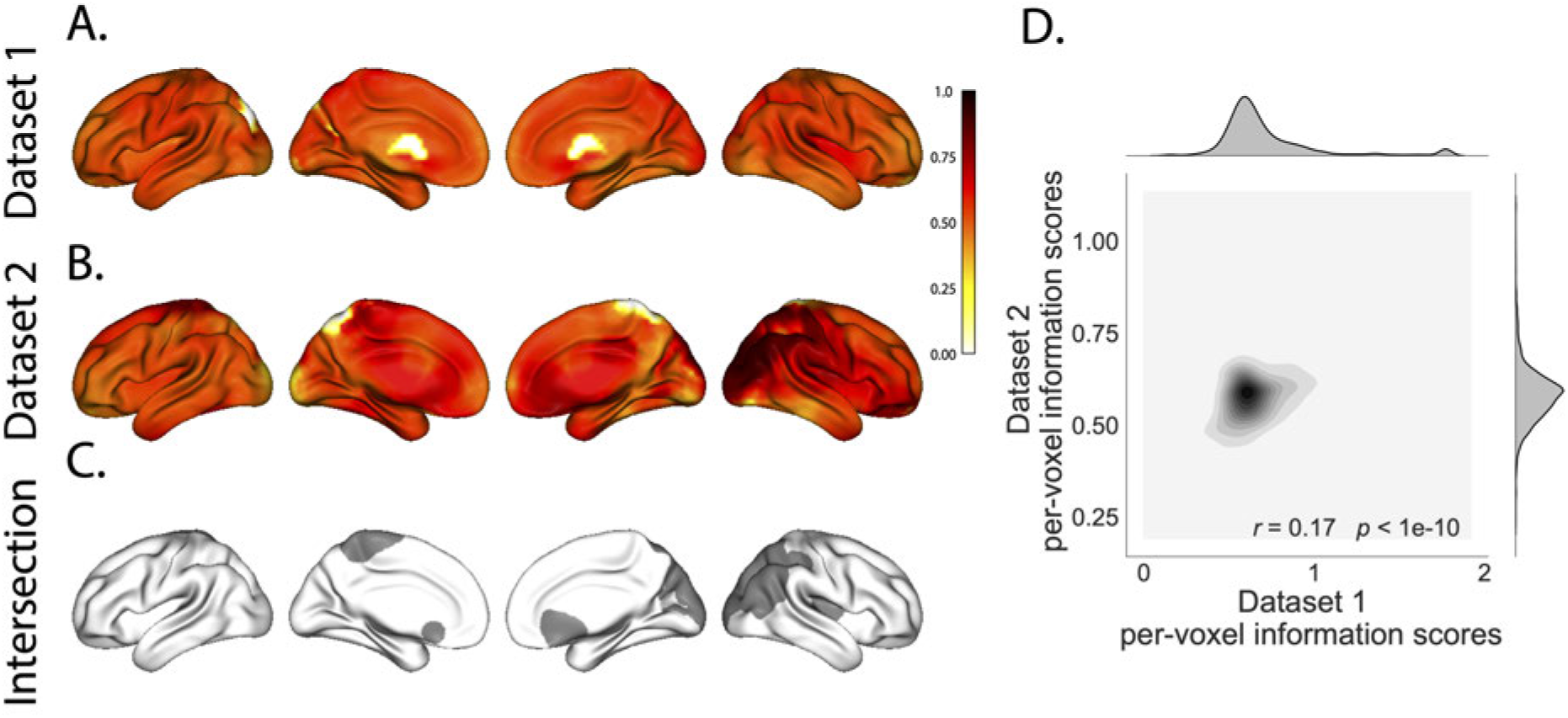
Most informative recording locations. **A. Dataset 1 information scores by voxel.** The voxel colors reflect the weighted average reconstruction accuracy across all electrodes from any patients with at least one electrode within 20 MNI units of the given voxel. **B. Dataset 2 information scores by voxel.** This panel is in the same format as Panel A. **C. Intersection.** Gray areas indicate the intersections between the top 10% most informative voxels in each map and projected onto the cortical surface (Combrisson et al. 2019). **D. Correspondence in information scores by voxel across Datasets 1 and 2.** The correlation reported in the Panel is between the per-voxel information scores across Datasets 1 and 2.

**Figure 6:**
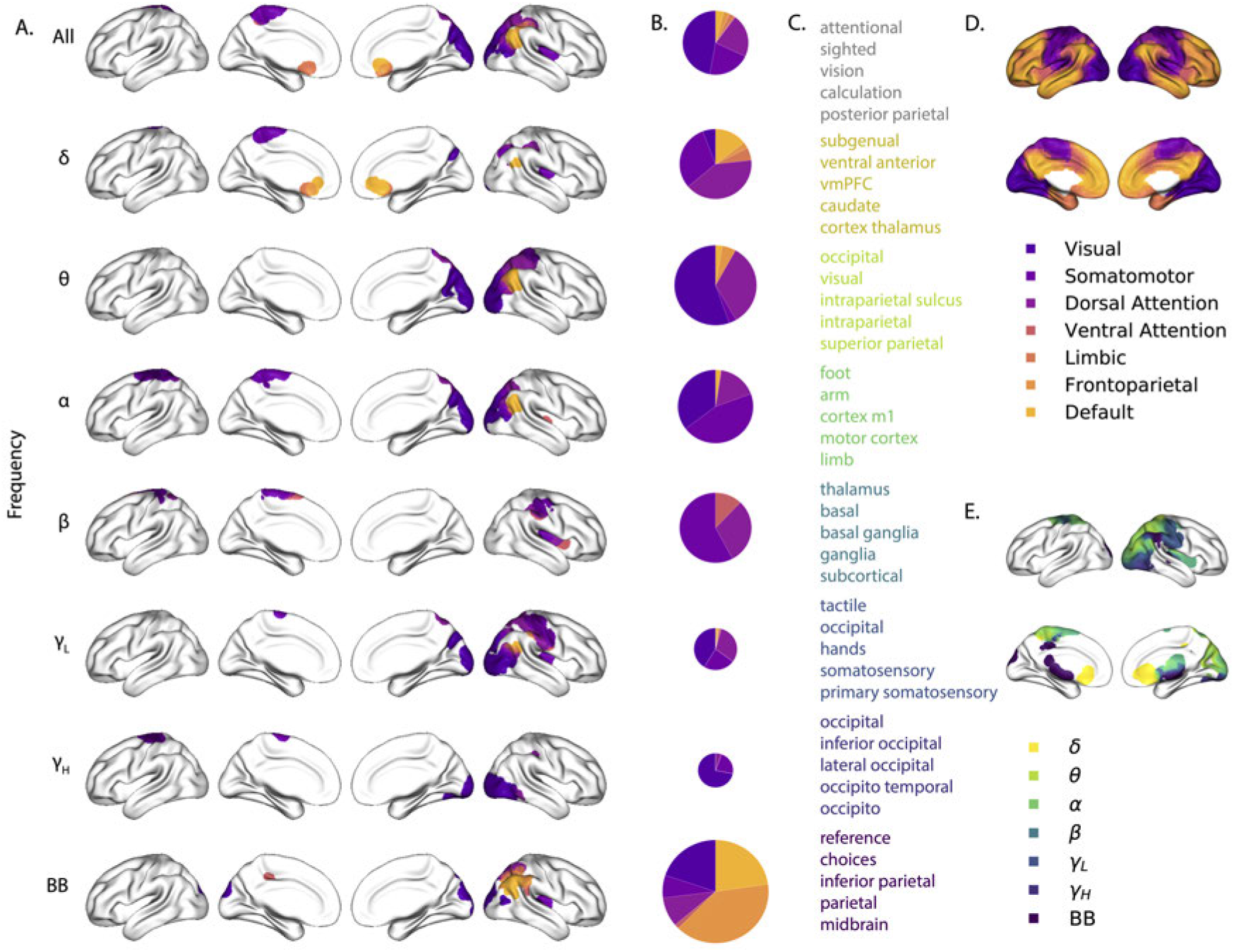
Most informative recording locations by frequency band. A. Intersections between information score maps by frequency band. The regions indicated in each row depict the intersection between the top 10% most informative locations across Datasets 1 and 2. **B. Network memberships of the most informative brain regions.** The pie charts display the proportions of voxels in each region that belong to the seven networks identified by Yeo et al. (2011). The relative sizes of the charts for each frequency band reflect the average across-subject reconstruction accuracies (Figs. 4A, D). The voxels in Panel A are colored according to the same network memberships. **C. Neurosynth terms associated with the most informative brain regions, by frequency band.** The lists in each row display the top five neurosynth terms (Rubin et al. 2017) decoded for each region. **D. Network parcellation map and legend.** The parcellation defined by Yeo et al. (2011) is displayed on the inflated brain maps. The colors and network labels serve as a legend for Panels A and B. **E. Combined map of the most informative brain regions.** The map displays the union of the most informative maps in Panel A, colored by frequency band. The labels also serve as a legend for Panel C.

We also wondered whether the map of information scores might vary as a function of the spectral components of the activity patterns under consideration. We computed analogous maps of information scores for each individual frequency band. Across Datasets 1 and 2 (with the exception of *α*-band activity), we observed reliable positive correlations between the voxelwise maps of information scores (*δ*: *r* = 0.09, *p* < 10^−57^; *θ*: *r* = 0.24, *p* < 10^−60^; *α*: *r* = −0.03, *p* < 0.001; *β*: *r* = 0.02, *p* = 0.0011; *γL* : *r* = 0.1, *p* < 10^−67^; *γH* : *r* = 0.03, *p* < 10^−7^; broadband: *r* = 0.21, *p* < 10^−297^).

To gain additional insight into which regions were most informative about full-brain activity patterns at different frequency bands, we next computed (for each frequency band) the intersection of the top 10% highest information scores across the maps for Datasets 1 and 2 (analogous to our approach in Fig. 5C). This yielded a single map of the (reliably) most informative locations, for each frequency band we examined. We then carried out *post hoc* analyses on each of these maps to characterize the underlying structural and functional properties of each set of regions we identified as being particularly informative about one or more types of neural pattern (Figs. 6 and S5).

A growing body of neuroscientific research is concerned with characterizing the *parcellations* of anatomical and functional brain networks (for review see Arslan et al. 2018; Zalesky et al. 2010). The dominant approaches entail obtaining a full-brain connectivity matrix using either diffusion tensor imaging to identify the brain’s network of white matter connections, or functional connectivity (typically applied to resting state data) to correlate the patterns of activity exhibited by different brain structures. One can then apply graph theoretic approaches to assign each brain structure (typically a single fMRI voxel) to one or more networks (for review see Bullmore and Sporns 2009). The result is a set of distinct (or partially overlapping) brain “networks” that may be further examined to elucidate their potential functional role. We overlaid a well-cited seven-network parcellation map identified by Yeo et al. (2011) onto the maps of brain locations that were most informative about each type of neural pattern. For each of these information maps, we computed the proportion of voxels in the most informative brain regions that belonged to each of the seven networks identified by Yeo et al. (2011); Figure 6D. We found that the regions we identified as being most informative about different neural patterns varied markedly with respect to which functional networks they belonged to (Fig. 6A, B).

The variability we observed in the frequency-specific information score maps is consistent with the notion that there is no “universal” brain region that reflects all types of activity patterns throughout the rest of the brain. Rather, each region’s activity patterns appear to be characterized by different spectral profiles, and the ability to infer full-brain activity patterns at a particular frequency band depends on the structural and functional connectome specific to that frequency band (Fig. 6E). We wondered how the maps we found might fit in with prior work. To this end, in addition to examining the anatomical profiles of each map, we used Neurosynth (Rubin et al. 2017) to identify (using meta analyses of the neuroimaging literature) the top five most common terms associated with each frequency-specific map (Fig. 6C). We found that *δ* patterns across the brain were best predicted by regions of ventromedial prefrontal cortex, striatum, and thalamus (yellow). These regions are also implicated in modulating *δ* oscillations during sleep, and are heavily interconnected with cortex (e.g., Amzica and Steriade 1998). The brain areas most informative about full-brain *θ* patterns were occipital and parietal regions associated with visual processing and visual attention (light green). Prior work has implicated *θ* oscillations in these areas in periodic sampling of visual attention (e.g., Busch and VanRullen 2010). We found that full-brain *α* patterns were best predicted by motor areas (dark green), which also exhibit *α* band changes during voluntary movements (e.g., Jurkiewicz et al. 2006). Striatum and thalamus (teal) were most informative about full-brain *β* patterns. Prior work has implicated striatal *β* activity in sensory and motor processing (Feingold et al. 2015) and thalamic *β* activity has been implicated in modulating widespread *β* patterns across neocortex (Sherman et al. 2016). Somatosensory areas (dark blue) were most informative about full-brain *γL* patterns. Prior work has implicated somatosensory *γL* in somatosensory processing and motor planning (Ihara et al. 2003). Occipital cortex (purple) was most informative about full-brain γ*H* patterns. Occipital γ*H* has also been linked with visual processing and reading (Wu et al. 2011) and the transmission of visual representations from low-order to higher-order visual areas (Matsumoto et al. 2013). Full-brain broadband patterns were best predicted by inferior parietal cortex precuneus (maroon). Functional neuroimaging BOLD responses (Simony et al. 2016) and broadband ECoG patterns (Honey et al. 2012) in these default-mode hubs have been implicated in processing context-dependent representations that unfold over long timescales.

Taken together, the frequency-specific information maps suggest a potential new interpretation of many of the above previously reported findings. Prior work has largely treated region-specific narrowband and broadband activity as an indicator that activity at those frequency ranges reflects that the given region is representing or supporting a particular function. Our work suggests an alternative interpretation that when we observe a particular neural pattern in a particular brain region, it may instead reflect how that region is transmitting information to the rest of the brain via signalling at the given frequency range.

## Discussion

Are our brain’s networks static or dynamic? And to what extent are the network properties of our brains stable across people and tasks? One body of work suggests that our brain’s *functional* networks are dynamic (e.g., Manning et al. 2018; Owen et al. 2019), person-specific (e.g., Finn et al. 2015), and task-specific (e.g., Turk-Browne 2013). In contrast, although the gross anatomical structure of our brains changes meaningfully over the course of years as our brains develop, on the timescales of typical neuroimaging experiments (i.e., hours to days) our anatomical networks are largely stable (e.g., Casey et al. 2000). Further, many aspects of brain anatomy, including white matter structure, are largely preserved across people (e.g., Jahanshad et al. 2013; Mori et al. 2008; Talairach and Tournoux 1988). There are several possible means of reconciling this apparent inconsistency between dynamic person- and task-specific functional networks versus stable anatomical networks. For example, relatively small magnitude anatomical differences across people may be reflected in reliable functional connectivity differences. Along these lines, one recent study found that diffusion tensor imaging (DTI) structural data is similar across people, but may be used to predict person-specific resting state functional connectivity data (Becker et al. 2018). Similarly, other work indicates that task-specific functional connectivity may be predicted by resting state functional connectivity data (Cole et al. 2016; Tavor et al. 2016). Another (potentially complementary) possibility is that our functional networks are constrained by anatomy, but nevertheless exhibit (potentially rapid) task-dependent changes (e.g., Sporns and Betzel 2016).

Here we have taken a model-based approach to studying whether high spatiotemporal resolution activity patterns throughout the human brain may be explained by a static connectome model that is shared across people and tasks. Specifically, we trained a model to take in recordings from a subset of brain locations, and then predicted activity patterns during the same interval, but at *other* locations that were held out from the model. Our model, based on Gaussian process regression, was built on three general hypotheses about the nature of the correlational structure of neural activity (each of which we tested). First, we hypothesized that functional correlations are stable over time and across tasks. We found that, although aspects of the patients’ functional correlations were stable across tasks, we achieved better reconstruction accuracy when we trained the model on within-task data. This suggests that our general approach could be extended to better model across-task changes, e.g., following Cole et al. (2016); Tavor et al. (2016); and others. Second, we hypothesized that some of the correlational structure of people’s brain activity is similar across individuals. Consistent with this hypothesis, our model explained each patient’s data best when trained using data from *other* patients– even when compared models trained within-patient. Third, we resolved ambiguities in the data by hypothesizing that neural activity from nearby sources tends to be similar, all else being equal. This hypothesis was supported through our finding that all of the models we trained that incorporated this spatial smoothness assumption predicted held-out data well above chance.

One potential limitation of our approach is that it does not provide a natural means of estimating the precise timing of single-neuron action potentials. Prior work has shown that gamma band and broadband activity in the LFP may be used to estimate the firing rates of neurons that underly the population contributing to the LFP (Crone et al. 2011; Jacobs et al. 2010; Manning et al. 2009; Miller et al. 2008). Because SuperEEG reconstructs LFPs throughout the brain, one could in principle use broadband power in the reconstructed signals to estimate the corresponding firing rates (though not the timings of individual action potentials). We found that we were able to reconstruct full-brain patterns of broadband power well (Fig. 4).

A second potential limitation of our approach is that it relies on ECoG data from epilepsy patients. Recent work comparing functional correlations in epilepsy patients (measured using ECoG) and healthy individuals (measured using fMRI) suggests that there are gross similarities between these populations (e.g., Kucyi et al. 2018; Reddy et al. 2018). Nevertheless, because all of the patients we examined have drug-resistant epilepsy, it remains uncertain how generally the findings reported here might apply more broadly to the population at large (e.g., non-clinical populations).

Beyond providing a means of estimating ongoing activity throughout the brain using already-implanted electrodes, our work also has implications how to optimize electrode placements in neurosurgical evaluations. Electrodes are typically implanted to maximize coverage of suspected epileptogenic tissue. However, our findings suggest that this approach might be improved upon. Specifically, one could leverage not only the non-invasive recordings taken during an initial monitoring period (as is currently done routinely), but also recordings collected from *other* patients. We could then ask: given what we learn from other patients’ data (and potentially from the scalp EEG recordings of this new patient), where should we place a fixed number of electrodes to maximize our ability to map seizure foci? As shown in Figures 5, 6, and S5, recordings from different regions vary with respect to how informative they are about different narrowband and broadband full-brain activity patterns.

By providing a means of reconstructing full-brain activity patterns, the SuperEEG approach maps ECoG recordings from different patients into a common neural space, despite that different patients’ electrodes were implanted in different locations. This feature of our approach enables across-patient ECoG studies, analogous to across-subject fMRI studies (e.g., Haxby et al. 2001, 2011; Norman et al. 2006). Whereas the focus of this manuscript is to specifically evaluate which aspects of neural activity patterns SuperEEG recovers well (or poorly), in parallel work we are training across-patient classifiers by leveraging the common neural spaces obtained by applying SuperEEG to multi-patient ECoG data. For example, we have shown that SuperEEG-derived activity patterns may be used to accurately predict psychiatric conditions such as depression (Scangos et al. 2020). Analogous approaches could in principle be used to develop improved brain-computer interfaces and/or to carry out other analyses that would benefit from high spatiotemporal resolution full-brain data in individuals, projected into a common ECoG space across people.

### Concluding remarks

Over the past several decades, neuroscientists have begun to leverage the strikingly profound mathematical structure underlying the brain’s complexity to infer how our brains carry out computations to support our thoughts, actions, and physiological processes. Whereas traditional beamforming techniques rely on geometric source-localization of signals measured at the scalp, here we propose an alternative approach that leverages the rich correlational structure of two large datasets of human intracranial recordings. In doing so, we are one step closer to observing, and perhaps someday understanding, the full spatiotemporal structure of human neural activity.

## Code availability

We have published an open-source toolbox implementing the SuperEEG algorithm. It may be downloaded <u>here</u>. Additionally, we have provided code for all analyses and figures reported in the current manuscript, available <u>here</u>.

## Data availability

The datasets analyzed in this study were generously shared by Michael J. Kahana. A portion of Dataset 1 may be downloaded <u>here</u>. Dataset 2 may be downloaded <u>here</u>.

## Acknowledgements

We are grateful for useful discussions with Luke J. Chang, Uri Hasson, Josh Jacobs, Michael J. Kahana, and Matthĳs van der Meer. We are also grateful to Michael J. Kahana for generously sharing the ECoG data we analyzed in our paper, which was collected under NIMH grant MH55687 and DARPA RAM Cooperative Agreement N66001-14-2-4-032, both to M.J.K. Our work was also supported in part by NSF EPSCoR Award Number 1632738 and by a sub-award of DARPA RAM Cooperative Agreement N66001-14-2-4-032 to J.R.M. The content is solely the responsibility of the authors and does not necessarily represent the official views of our supporting organizations.

## Author Contributions

J.R.M conceived and initiated the project. L.L.W.O., T.A.M, and A.C.H. performed the analyses using software packages that all authors contributed to. J.R.M. and L.L.W.O. wrote the manuscript with input from all other authors.

## Footnotes

1 The term “SuperEEG” was coined by Robert J. Sawyer in his popular science fiction novel *The Terminal Experiment* (Sawyer 1995). SuperEEG is a fictional technology that measures ongoing neural activity throughout the entire living human brain at arbitrarily high spatiotemporal resolution.

