## Supporting Information for "A Gaussian process model of human electrocorticographic data"

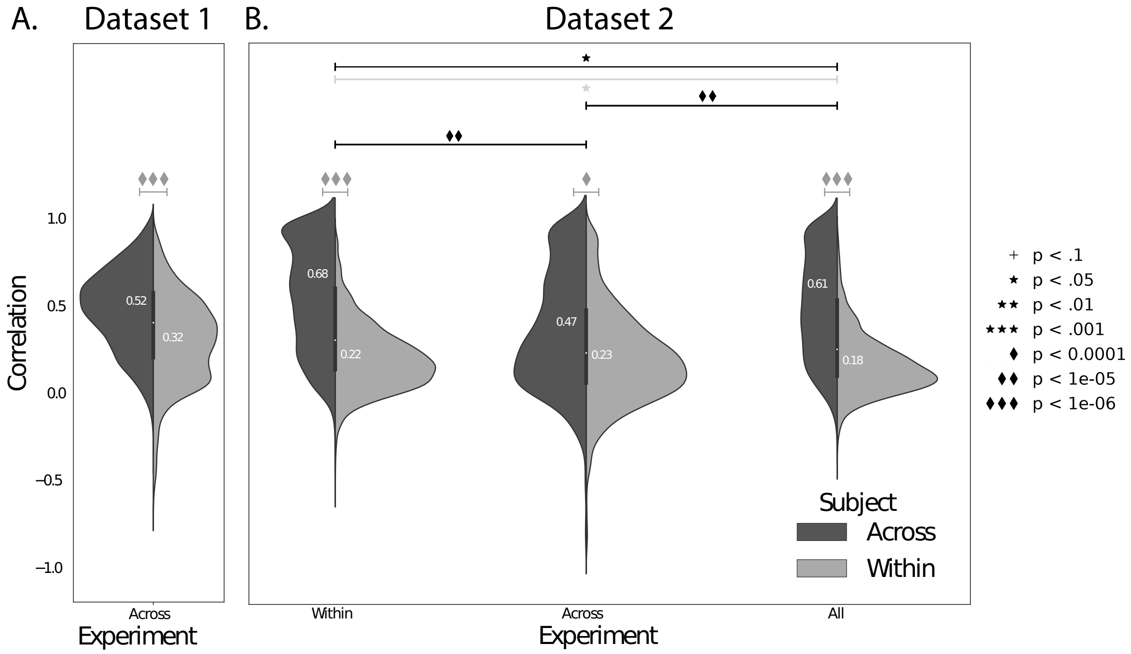


**Figure S1: Reconstruction accuracy across all electrodes in two ECoG datasets, broken down by experiment. A. Distributions of correlations between observed versus reconstructed activity by electrode, for Dataset 1.** The across-patient distribution (black) reflects reconstruction accuracy (correlation) using a correlation model learned from all but one patient's data, and then applied to that held-out patient's data. The within-patient distribution (gray) reflects performance using a correlation model learned from the same patient who contributed the to-be-reconstructed electrode. The split violin plot displays the same distributions shown in Figure 2A. The white numbers within each half denote the means of each distribution. **B. Distributions of correlations for Dataset 2.** The split violin plots are in the same format as those in Panel A. The leftmost plot ("Within") displays the same distributions shown in Figure 2B. All reconstructions reflected in the distribution were carried out using a model trained and tested using data from the same experiment. The middle plots ("Across") reflect reconstructions trained and tested using data from different experiments. The rightmost plot ("All") reflect reconstructions obtained using models trained and tested on data from both experiments. The black distributions reflect models trained and tested across patients (analogous to the black histograms in Figure 2) and the gray distributions reflect models trained and tested within patient (analogous to the gray histograms in Figure 2). The dark gray significance bars reflect paired-sample $t$-tests comparing the ($z$-transformed) reconstruction accuracy for each electrode obtained within versus across patients. The black significance bars reflect paired-sample $t$-tests comparing the across-patient reconstruction accuracies across datasets. The light gray significance bars reflect paired-sample $t$-tests comparing the within-patient reconstruction accuracies across datasets. The symbols denote the corresponding $p$-values of those statistical tests.

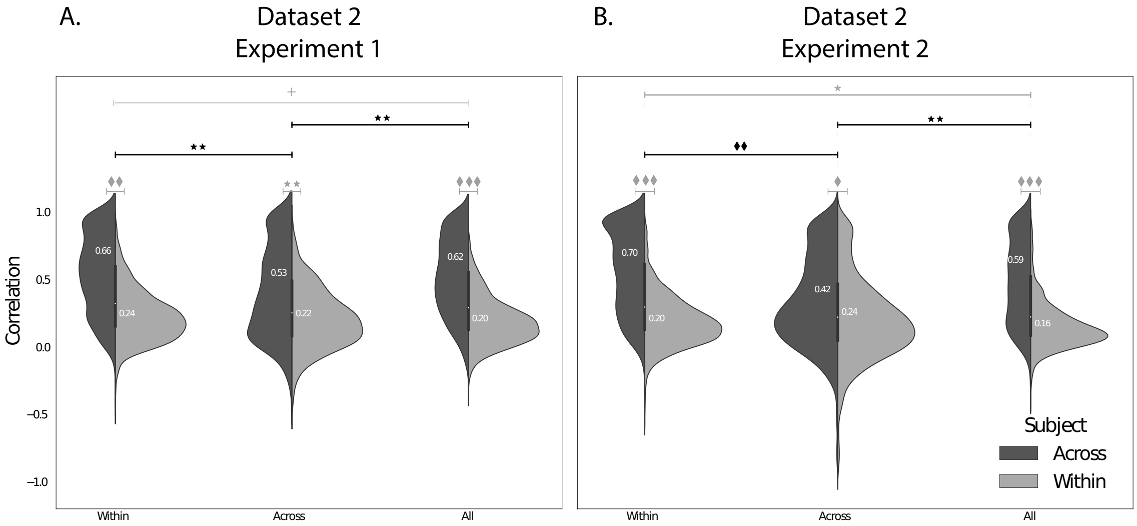


**Figure S2: Reconstruction accuracy for Dataset 2, Experiments 1 and 2. A. Distributions of correlations between observed versus reconstructed activity by electrode, for Dataset 2 (Experiment 1).** The plots are in the same format as Figure S1B, but reflect data only from Experiment 1 in Dataset 2. **A. Distributions of correlations between observed versus reconstructed activity by electrode, for Dataset 2 (Experiment 2).** The plots are in the same format as Panel A, but reflect data only from Experiment 2 in Dataset 2.

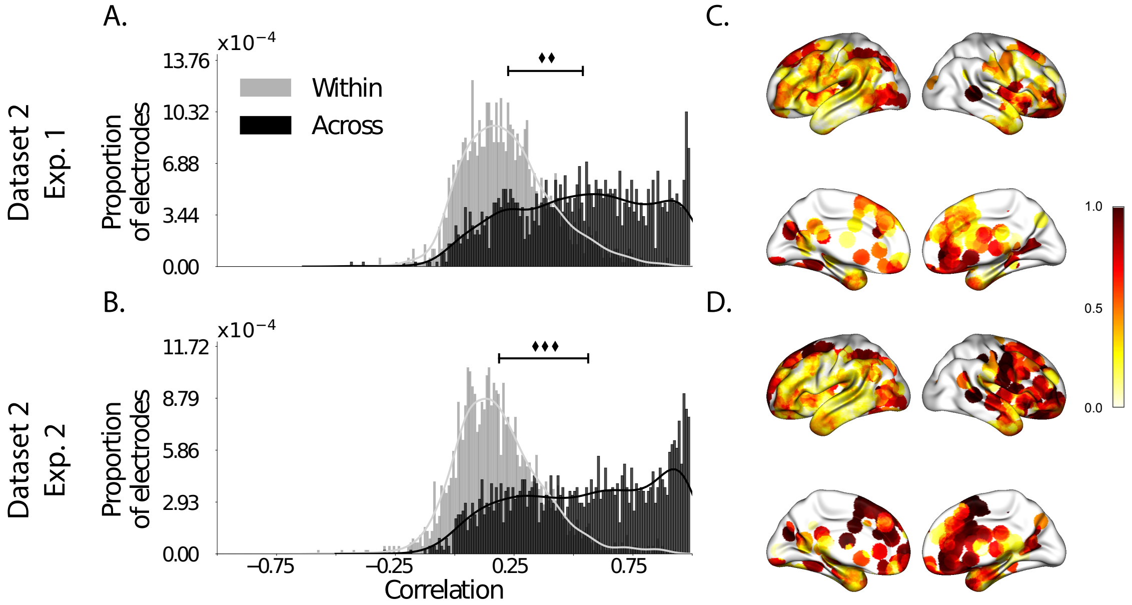


**Figure S3: Reconstruction accuracy across all electrodes in Dataset 2.** This figure is analogous to Figure 2 in the main text, but displays data for Dataset 2 broken down by experiment. **A. Distributions of correlations between observed versus reconstructed activity by electrode for Experiment 1.** The across-patient distribution (black) reflects reconstruction accuracy (correlation) using a correlation model learned from all but one patient's data, and then applied to that held-out patient's data. The within-patient distribution (gray) reflects performance using a correlation model learned from the same patient who contributed the to-be-reconstructed electrode. **B. Distributions of correlations for Experiment 2.** This panel is in the same format as Panel A, but reflects results obtained from Experiment 2. **C.–D. Reconstruction accuracy by location.** The colors denote the average across-session correlations, using the across-patient correlation model, between the observed and reconstructed activity at the given electrode location projected to the cortical surface (Combrisson et al., 2019). Panel C displays the map for Experiment 1 and Panel D displays the map for Experiment 2.

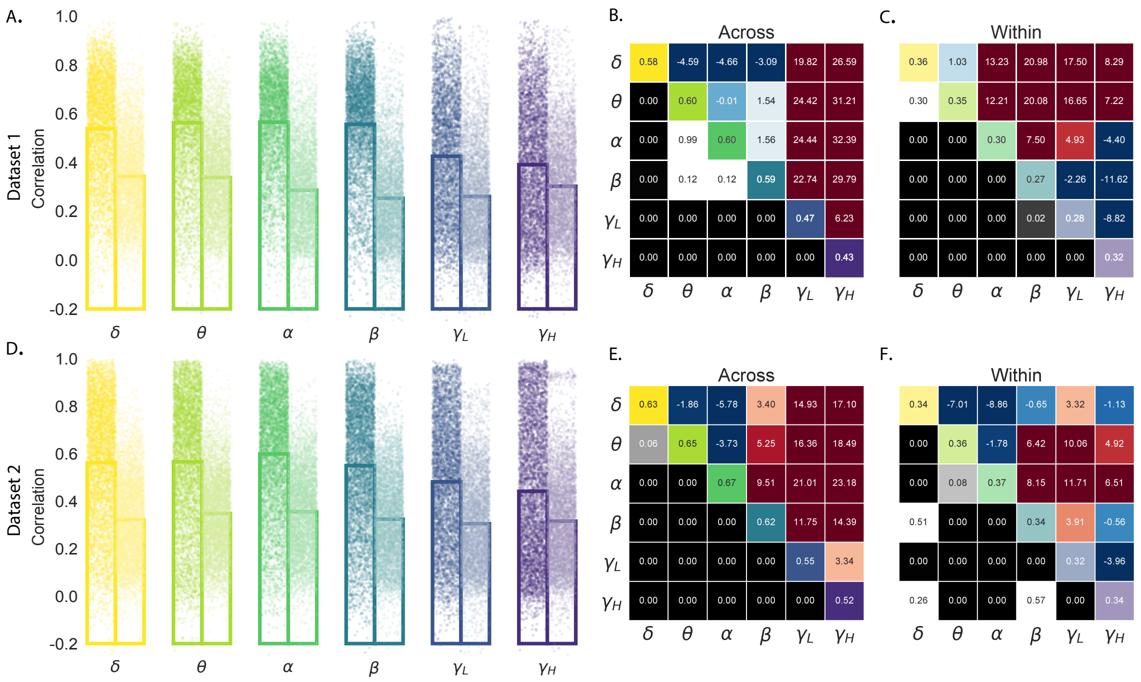


**Figure S4: Reconstruction accuracy across all electrodes in two ECoG datasets for power in each frequency band.** This figure is analogous to Figure 3 in the main text, but displays the reconstruction accuracies of the *power* in each frequency band, rather than the bandpass-filtered voltages, as in Figure 3. **A. Distributions of correlations between observed versus reconstructed power by electrode for each frequency band in Dataset 1.** Each color denotes a different frequency band. Within each color group, the darker dots and bar on the left display the distribution (and mean) across-patient reconstruction accuracies (analogous to the black histograms in Fig. 2). The lighter dots and bar on the right display the distribution (and mean) within-patient reconstruction accuracies (analogous to the gray histograms in Fig. 2). Each dot indicates the reconstruction accuracy for one electrode in the dataset. **B. Statistical summary of across-patient reconstruction accuracy by electrode for each frequency band in Dataset 1.** In the upper triangles of each map, warmer colors (positive *t*-values) indicate that the reconstruction accuracy for the frequency band in the given row was greater (via a two-tailed paired-sample *t*-test) than for the frequency band in the given column. Cooler colors (negative *t*-values) indicate that reconstruction accuracy for the frequency band in the given row was lower than for the frequency band in the given column. The lower triangles of each map denote the corresponding *p*-values for the *t*-tests. The diagonal entries display the average reconstruction accuracy within each frequency band. **C. Statistical summary of within-patient reconstruction accuracy by electrode for each frequency band in Dataset 1.** This panel displays the within-patient statistical summary, in the same format as Panel B. **D. Distributions of correlations between observed versus reconstructed activity by electrode, for each frequency band in Dataset 2.** This panel displays reconstruction accuracy distributions for each frequency band for Dataset 2. **E.–F. Statistical summaries of across-patient and within-patient reconstruction accuracy by electrode for each frequency band in Dataset 2.** These panels are in the same as Panels B and C, but display results from Dataset 2.

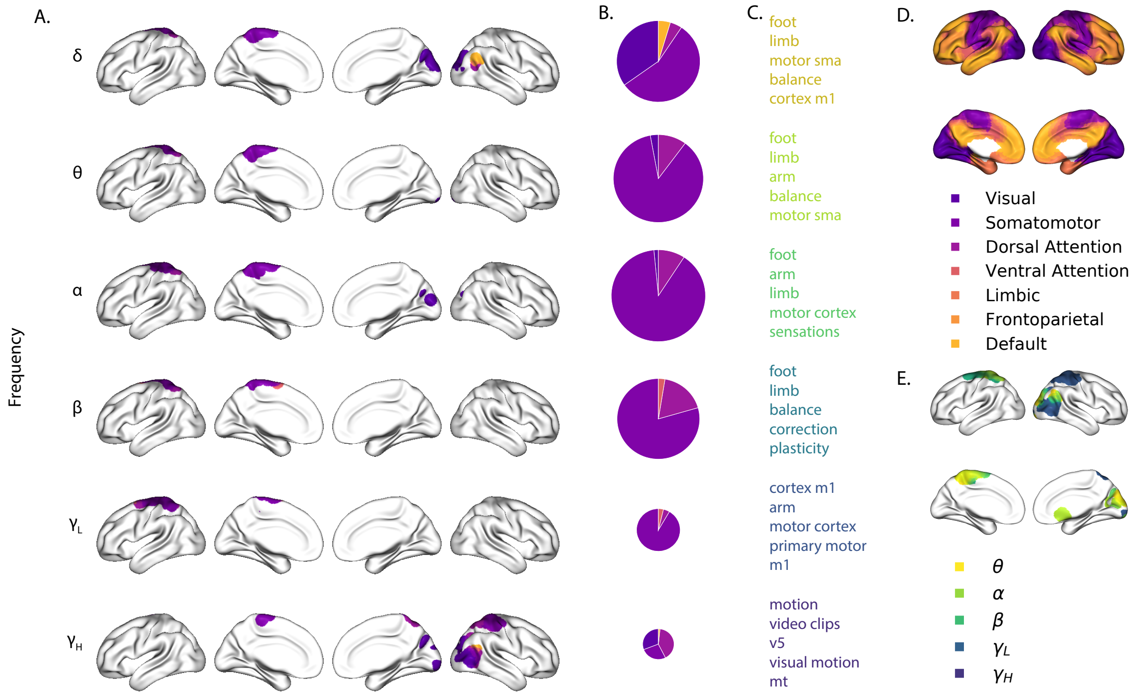


**Figure S5: Most informative recording locations by frequency band.** This figure is analogous to Figure 6 in the main text, but displays the brain regions that were most informative about the *power* in each frequency band (whereas Fig. 6 displays the regions that were most informative about the bandpass-filtered voltages within each band). **A. Intersections between information score maps by frequency band.** The regions indicated in each row depict the intersection between the top 10% most informative locations across Datasets 1 and 2. **B. Network memberships of the most informative brain regions.** The pie charts display the proportions of voxels in each region that belong to the seven networks identified by Yeo et al. (2011). The relative sizes of the charts for each frequency band reflect the average across-subject reconstruction accuracies (Figs. 3A, D). The voxels in Panel A are colored according to the same network memberships. **C. Neurosynth terms associated with the most informative brain regions, by frequency band.** The lists in each row display the top five neurosynth terms (Rubin et al., 2017) decoded for each region. **D. Network parcellation map and legend.** The parcellation defined by Yeo et al. (2011) is displayed on the inflated brain maps. The colors and network labels serve as a legend for Panels A and B. **E. Combined map of the most informative brain regions.** The map displays the union of the most informative maps in Panel A, colored by frequency band. The labels also serve as a legend for Panel C.

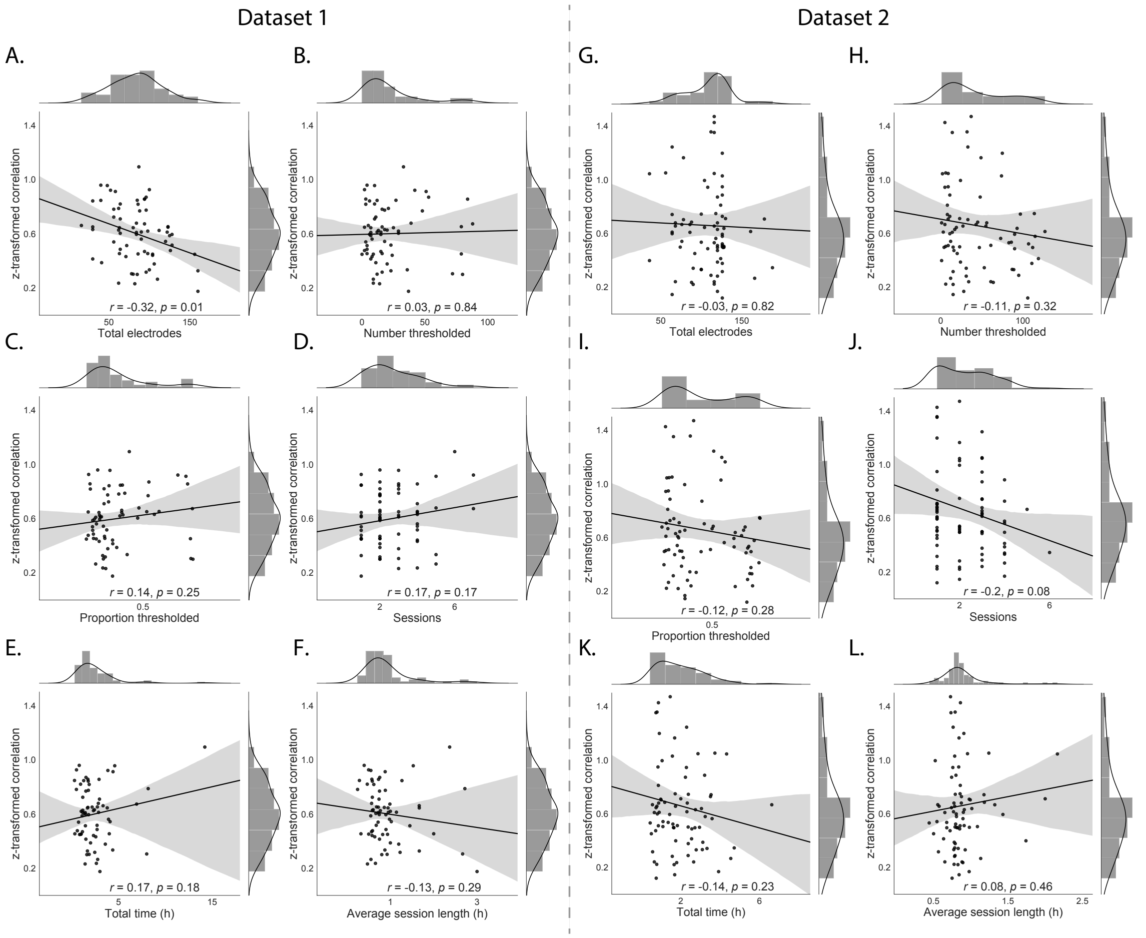


**Figure S6: Reconstruction accuracy versus within-subject data features for two ECoG datasets.** The individual dots in each panel reflect data from one patient. The least-squares linear regression lines (with shaded 95% confidence intervals) and the correlations reported in each panel are between the data shown on the $x$- and $y$-axes of the panel. The $y$-axis in each panel denotes the average $z$-transformed correlation between the observed and predicted voltage timeseries at each electrode (across all of the patient's electrodes). The cross-validated predictions were obtained using the across-patient model (Fig. 2A and B, black histograms). **A.–F. Dataset 1. A. Total electrodes.** The $x$-coordinates of each dot display the total number of electrodes implanted in each patient's brain. **B. Number thresholded.** The $x$-coordinates of each dot display the number of implanted electrodes that survived the kurtosis-based filtering procedure (see *Approach*). **C. Proportion thresholded.** The $x$-coordinates of each dot display the *proportion* of each patient's implanted electrodes (relative to the total number) that survived the kurtosis-based filtering procedure. **D. Sessions.** The $x$-coordinates of each dot display the number of distinct recording sessions contributed by each patient. **E. Total time.** The $x$-coordinates of each dot display the total duration (in hours) of each patient's recordings, across all of their recording sessions. **F. Average session length.** The $x$-coordinates of each dot display average duration (in hours) of the patient's recording sessions. **G.–L. Features from Dataset 2.** These panels display analogous information to Panels A–F, but for Dataset 2 patients.

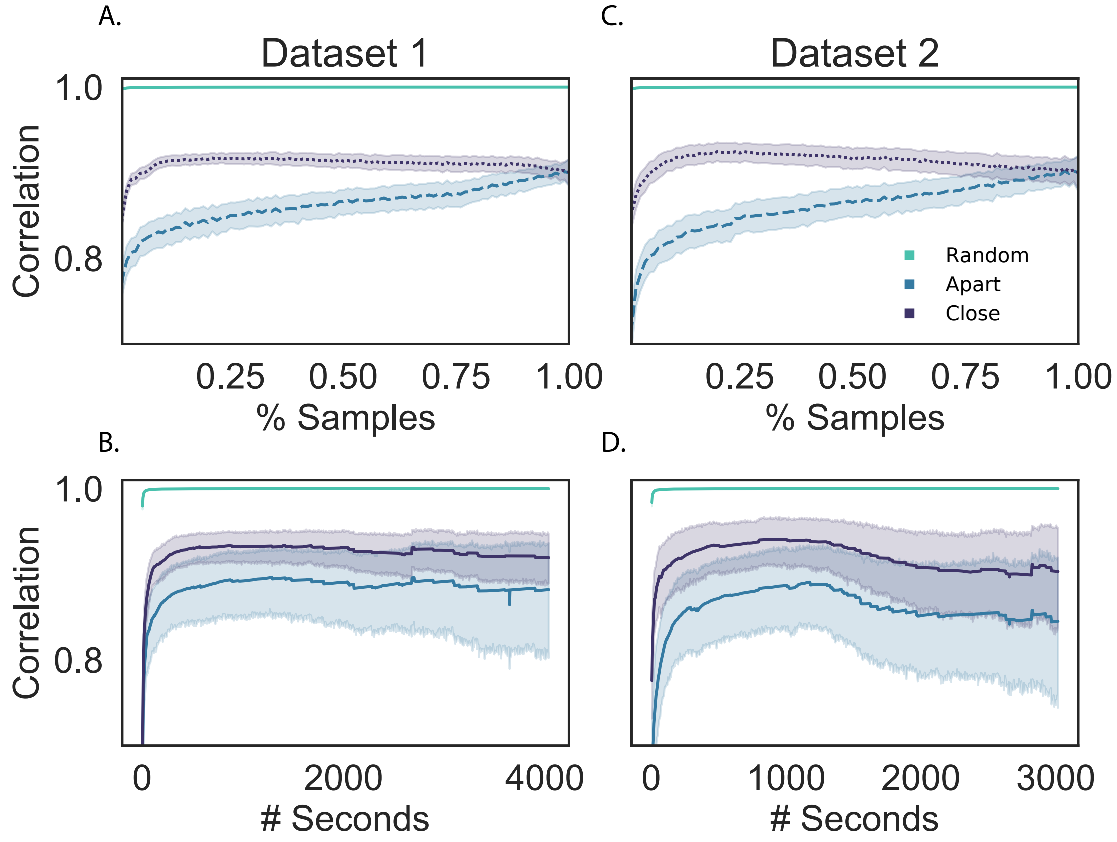


**Figure S7: Temporal stability of estimated patient-specific correlation matrices.** For each patient $s$, we computed their patient-specific correlation matrix ($C_s$) using two different subsets of their recorded data. We correlated the upper triangles of these estimates of $C_s$ to obtain a single correlation for each participant, reflecting how stable (in time) the estimates were. We computed this stability measure, for each patient, as a function of how much data was used to estimate each $C_s$ (expressed as a proportion of the patient's total dataset, as in Panels **A** and **C**; or as the total duration of the patient's recordings, as in Panels **B** and **D**). We also varied how the timepoints that went into each estimate were related. Specifically, we first concatenated the data from each patient's recording sessions to construct a single data timeseries (ignoring session boundaries) for each patient. We then constructed two estimates of $C_s$ by drawing each patient's data from two equally sized sets of $t$ timepoints chosen from their multi-session data matrix. These timepoints were drawn either at *random* without replacement (teal lines); from equally sized timespans at the start and end of the multi-session data matrix (*apart*, blue lines); or from two equally sized timespans just prior to and just after the data midpoint (*close*, purple lines). For the random condition of this analysis, as we increased $t$, we ensured that all of the timepoints included in the analysis for smaller values of $t$ were also included in the analyses for larger values of $t$. For example, the randomly drawn timepoints for $t = 2000$ included the same timepoints drawn for $t = 1000$, plus an additional 1000 new timepoints. Panels A and B display the resulting correlations for Dataset 1, and panels C and D display the correlations for Dataset 2. Error ribbons in all panels denote 95% confidence intervals (computed across patients).

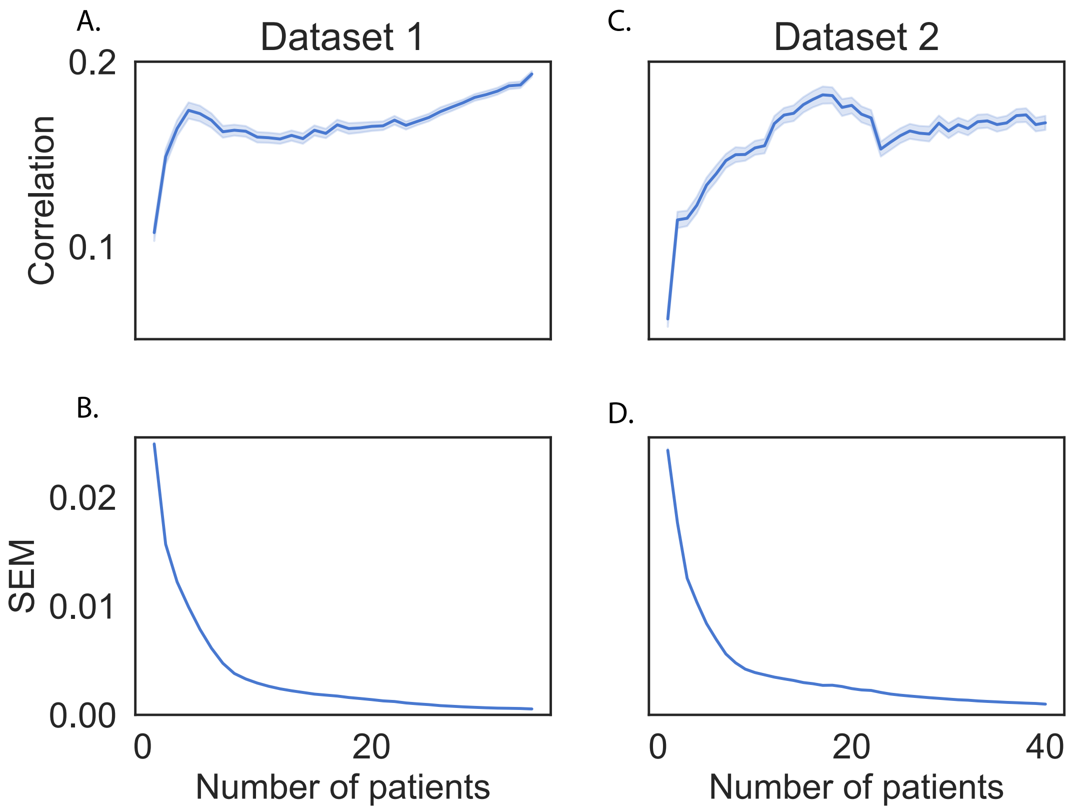


**Figure S8: Stability of full-brain correlation matrices across patients.** We estimated full-brain correlation matrices ($\hat{K}$) using different subsets of patients from each dataset. We explored how stable these estimates were as a function of how many patients were used to compute $\hat{K}$. For each sample size $n$ (number of patients), we drew two non-overlapping sets of $n$ patients (without replacement) from the full set of patients in each dataset. We then estimated $\hat{K}$ using the two sets of $n$ patients. We computed the correlation between the upper triangles of these matrices as a measure of how similar the matrices were. We repeated this procedure 500 times for each value of $n$ (ranging from 1 up to $\frac{1}{2}$ of the total number of patients in the given dataset); this yielded a distribution of correlation coefficients for each value of $n$. Panels **A** and **C** display the mean correlations as a function of $n$ for Datasets 1 and 2, respectively. The error ribbons denote 95% confidence intervals (computed across iterations). Panels **B** and **D** display the standard error of the mean (SEM) of these distributions of correlations as a function of $n$, for Datasets 1 and 2, respectively. The error ribbons denote bootstrap-derived 95% confidence intervals.
